# Low-Bias Amplification for Robust DNA Data Readout

**DOI:** 10.1101/2020.02.09.940411

**Authors:** Yanmin Gao, Xin Chen, Jianye Hao, Chengwei Zhang, Hongyan Qiao, Yonggang Ke, Hao Qi

## Abstract

In DNA data storage, the massive sequence complexity creates challenges in repeatable and efficient information readout. Here, our study clearly demonstrated that canonical polymerase chain reaction (PCR) created significant DNA amplification biases, which greatly hinder fast and stable data retrieving from hundred-thousand synthetic DNA sequences encoding over 2.85 megabyte (MB) digital data. To mitigate the amplification bias, we adapted an isothermal DNA amplification for low-bias amplification of DNA pool with massive sequence complexity, and named the new method isothermal DNA reading (iDR). By using iDR, we were able to robustly and repeatedly retrieve the data stored in DNA strands attached on magnetic beads (MB) with significantly decreased sequencing reads, compared with the PCR method. Therefore, we believe that the low-bias iDR method provides an ideal platform for robust DNA data storage, and fast and reliable data readout.

## Introduction

DNA, the primary molecular carrier of genetic information in biology, has attracted great interests in high-density, stable data storage recently. Important technologies, including array oligo synthesis, PCR, and DNA sequencing, originated from other research fields, have been integrated into DNA-based information storage(*1–4*). For example, chip-synthesized DNA oligo pool comprising of up to millions of oligo strands, each is several hundred bases in length, has been utilized in many applications, e.g., probe blend, DNA origami assembly, and genome synthesis(*1*). Due to the synthesis biases, substantial variation exists in the copy numbers of different DNA strands produced on microchip(*5*). Furthermore, the amount and concentration of each DNA sequence in an array on microchip is very low, roughly in the range of 10^5^ to 10^12^ copies and femtomolar concentration, depending on the synthesis platforms(*6, 7*). However, over a few hundred nanogram DNA, at micromolar concentration, is generally necessary for reliable information readout via commercial DNA sequencing platform, e.g., Illumina(*8*). Therefore, amplification of the chip-synthesized DNA is crucial to boost the signals for the subsequent DNA sequencing for robust retrieving of data.

The oligo pool sizes for DNA data storage are at least several orders of magnitude larger than those used in many biomedical applications(*9–11*). The smallness and the unevenness of DNA copy numbers cause a serious problem for DNA sequence retrieval and data decoding(*5, 12, 13*). Thus far, the information retrieving in reported systems was achieved by PCR amplification and next-generation sequencing (NGS). However, the unevenness of copy numbers originally stemmed from the microchip synthesis was further skewed by highly biased PCR amplification process(*5, 12*). The length and sequence context, GC content and secondary structure of DNA molecule are well known to introduce large amplification bias in PCR for amplification of multiple templates in parallel(*14*). Oligos of low copy numbers could be easily excluded in a few amplification cycles because of the replication biases. Therefore, a large amount of DNA and a high number of sequencing reads were necessary to fetch the scarce oligo strands from the skewed oligo pool. Thus far, this problem was dealt by high encoding, physical and sequencing redundancy(*13, 15*). However, high redundancy sacrifices storage density and increases cost in synthesis, sequencing and decoding calculation, especially in large-scale DNA storage that requires amplification and sequencing of massive numbers of DNA sequences at the same time

In previous studies, it has been demonstrated that deep PCR amplification (over 60 thermal amplification cycles) significantly increased the unevenness of copy numbers and the subsequent decoding required several orders of magnitude more sequencing reads (*12*). In extreme cases, only a small portion of oligos (less than 10 % of the total oligos) wiped out almost all others after amplification(*16*). Another crucial issue for repeated PCR amplification is the thermal cycling process, which not only consumes energy but also causes complication for the operation of storage devices. It has been reported that long thermal treatment at around 65°C resulted to oligo decay, especially to longer DNA molecules(*16, 17*). Therefore, for more practical DNA data storage, it is necessary to develop novel methods that provides low-bias amplification of large oligo pools for reliable information retrieving and also supports repeated reading for stable long-term information storage. We employed an isothermal DNA reading (iDR) method to achieve low-bias, low-error DNA amplification of oligo pools with high sequence complexity for robust DNA storage. We demonstrated that the iDR can significantly reduce the replication bias and error-spreading and oligo dropout. Moreover, we showed that a DNA pool immobilized on magnetic beads (MB) combined with isothermal reading, iDR method, can be used a highly efficient system for repeatedly readable, sustainable information storage. To the best of our knowledge, this is the first sustainable DNA storage system successfully achieved the stable information “deep reading”.

## Results

### BASIC code for DNA mediated distributed storage

In current DNA storage, the information in a digital file was divided and written into a large group of small DNA oligos. Every single oligo functions as an individual information carrying molecule. The information can be reconstituted by sequencing all the oligos (Fig. 1A). In this work, we adapted a BASIC code system(*18, 19*) recently developed for digital distributed file system for DNA storage, in which a high information density was achieved with a low encoding redundancy of 1.56%. Generally, the encoding process started by dividing the target file into non-overlapping groups and the split information was encoded into DNA sequence following an optimized encoding process (fig. S1 and Section S1) with two adjustable parameters K (corresponding to oligo number in one non-overlapping group, K was set as an constant value of 256) and L (corresponding to the length of oligo). Besides the normal digital file types, we also included genome sequence including human mitochondrion and one artificial bacteria cell (Section S2) with a different encoding strategy. Instead of directly saving genome information, the genome sequences were converted to binary files with a simple coding of T to 00, G to 01, C to 10 and A to 11. Through this encoding process, the genome sequences were rewritten into new nucleotide sequences, which allows us to avoid any complicated natural genome sequences that can cause difficulties in sequencing. An oligo pool was designed to store 2.85 MB files in the sequences of total 109,568 oligo strands with different lengths (Fig. 1A, fig. S2A and Table S1). Reed-Solomon code was used for error correction and coding redundancy. The DNA pool has a relative high information density of 1.65 bits/nt and a 1.56% coding redundancy, calculated by a previous reported method without modification (Section S3)(*13*). Successful decoding could be achieved as long as oligo dropout rate is less than 1.56%, corresponding to randomly losing 4 strands out of 256 (fig. S1B). Technically, higher encoding redundancy can tolerate higher dropout rates and successful decoding can be achieved from fewer sequencing reads. However, it is impractical to increase encoding redundancy, considering the synthesis cost is already 90% of the total cost of DNA storage in our system.

**Fig. 1.**
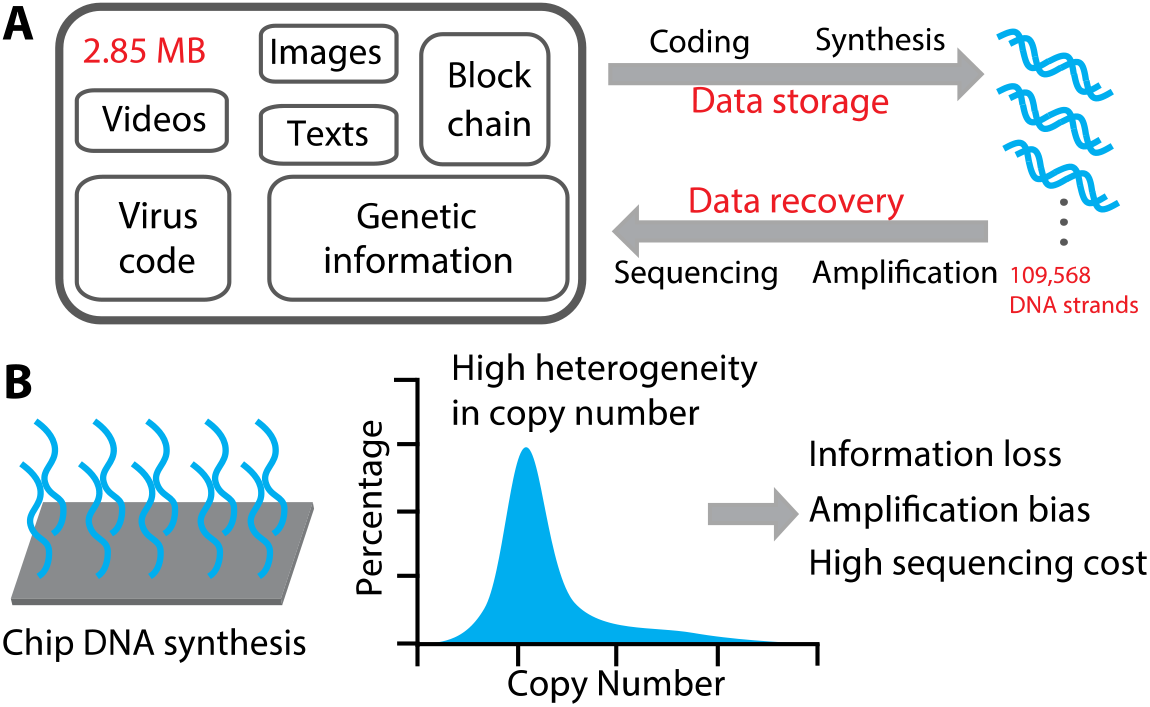
Overview of the DNA data storage. **(A)** Schematic of the DNA data storage and totally 2.85 MB data, including texts, images, videos, and block chain file, computer virus code, genetic information, were encoded into total 109,568 synthesis oligos using an adapted BASIC code system. **(B)** Illustration of data storage in synthesis oligo pool. The huge sequence complexity and unevenness of oligo copy number is challenging the stable storage and precise decoding.

Chip-based synthesis only produces very small amount of oligo, roughly from 10^5^ to 10^12^ copies at femtomolar concentrations(*7*). In the current workflow of DNA data storage, the quality of oligo pool greatly impacted the storage performance(*20*). Particularly, both the heterogenous oligo sequences and the unevenness of oligo copy number generated a huge sequence complexity and then caused the PCR amplification bias. Oligo molecules with low copy numbers are prone to drop-out and more sequencing coverage was required for decoding oligo pool with largely skewed copy number distribution (Fig. 1B).

### Low-bias and error-spreading proof isothermal amplification

It is well known that PCR generated biased amplification due to its mechanism, i.e., product-as-template, priming, and thermal cycling dependent amplification (Fig. 2A) (*3, 14, 21, 22*). To address these problems, we recruited isothermal DNA replications, which are majorly used in systems for biomolecular analysis or complex DNA circuit network (*23–25*). To the best of our knowledge, strand displacement replication has not been tested for robust amplification of large DNA libraries for DNA storage. Therefore, we decide to investigate whether we can design and build a low biased isothermal amplification of DNA pool with high sequence complexity. After systematical optimization of reaction temperature, reagent component, replicon length and nickase recognition site sequence (figs. S2B-E), Nt.BbvCI nickase and exonuclease deficient DNAP I Klenow fragment (3’→5’ exo^-^) were used and the amplification efficiency of iDR is around 70% of PCR (fig. S2F). The immobilized isothermal DNA replication system was designated as iDR, isothermal DNA reading for large DNA pool amplification (Fig. 2B). The Nt.BbvCI nickase recognition site is only 7 nts in length and its sequence has been avoided from the payload part of oligo pool in the file encoding step.

**Fig. 2.**
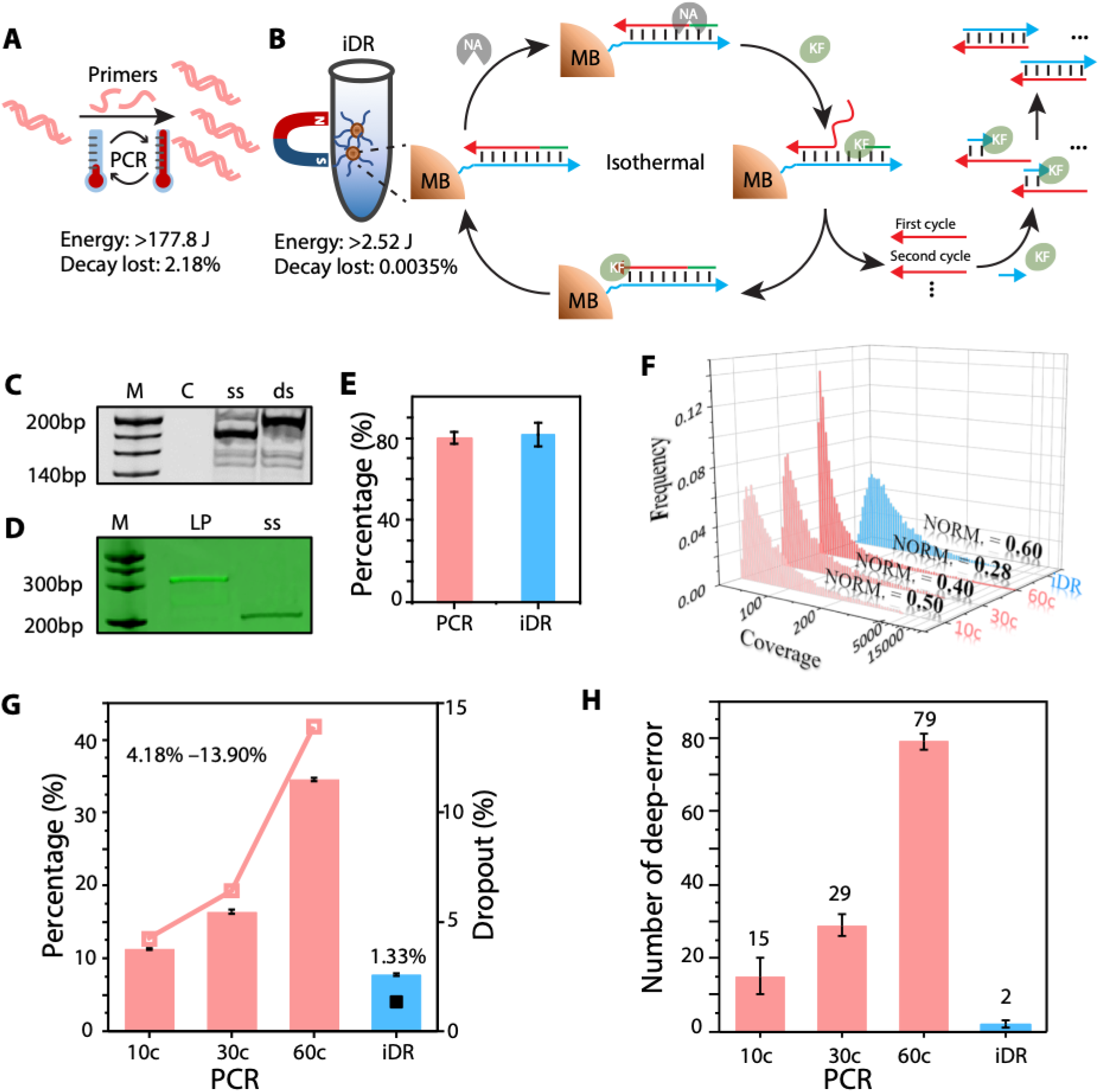
Designed isothermal DNA reading (iDR) for DNA data storage. **(A)** Illustration of the DNA amplification of PCR depending on thermal cycling to drive replication. For one 50 ul liquid reaction of 10 thermal cycles 177.8 J was required only for thermal regulation and caused 21.8‰ oligo decay theoretically. **(B)** Isothermal DNA reading (iDR) was developed from systematically optimized strand displacement mediated replication, in which nickase and specific DNA polymerase collaborate to drive amplification under consistent low temperature and only consume 2.52 J and 0.035‰ oligo decay due to thermal treatment. NA: nickase; KF: Klenow fragments (3’→5’ exo^-^). **(C)** Single-stranded and double-stranded oligo amplified by iDR from a 218nt dsDNA template were analyzed on 10% native PAGE gel. M: 20 bp DNA Ladder; C: negative control of amplification without input template; ss: single-stranded DNA product; ds: double-stranded DNA product. **(D)** A 5’ terminal FAM labeled 30 nt single-stranded probe ligated to 198 nt single-stranded DNA with an inborn 5’ terminal phosphate group directly from iDR amplification was analyzed and imaged on a 12% urea denaturing PAGE. M: 20 bp DNA Ladder; LP: ligation product; ss: ssDNA product of iDR. **(E)** Sequenced reads with single letter error, substitution or indel, accounted for the vast majority in total mutant sequenced reads for both PCR (80.1%) and iDR (81.7%). Error bars represent the mean ±s.d., where n = 3. **(F)** Coverage depth distribution of sequenced reads for 10, 30 and 60 cycles PCR and iDR amplification were positively skewed but with 0 oligo dropout. The distribution normality was 0.50 (10c), 0.40 (30c), 0.28 (60c), 0.60 (iDR) respectively. **(G)** The portion of oligo for the top 30% of high coverage which comprising 207 oligos for PCR sample and 197 for iDR. The reads of deep oligos increased significantly from 11.31% to 34.52% as the number of PCR cycles increased from 10 to 60. Meanwhile in iDR the high coverage reads only accounted for 7.77% in per million sequenced reads. Oligo dropout rate calculated from random reads set with a mean 10x coverage depth was plotted. For 10, 30 and 60 thermal cycles PCR (red line), it was counted as 4.18%, 6.41% and 13.90% respectively, and 1.33% for iDR (blue dot). Error bars represent the mean ±s.d., where n = 10. **(H)** The number of reads with mutant in a high copy number. The number increased significantly from 15±5 to 79±2 as the number of PCR cycles increased from 10 to 60. The number of iDR was 2±1.

Some intrinsic features of iDR makes it specifically suitable for DNA storage application. In theory, for one 50 ul reaction, 10 PCR thermal cycles required about 177.8J energy just for the thermal regulation, at least two orders of magnitude more than the 2.52J energy for 30 mins iDR (Fig. 2 A and B and Section S4). The larger energy consumption of PCR can be an issue for large-scale DNA storage. Real time monitoring indicated that 30 mins is the optimal amplification time (fig. S2G) and was used for all experiments in this manuscript. Single-stranded or double-stranded DNA can be produced in a controlled manner (Fig. 2C). Noticeably, ssDNA was amplified in a primer-free manner and allows iDR to be a universal method for reading information with no sequence information needed in advance for primer design. A nickase mediated site-specific phosphodiester bond cleavage initiated the iDR amplification and generated a 5’ terminal phosphate group, its function was verified by direct ligation to a FAM labeled probe (Fig. 2D). This phosphate group can be used for subsequent modification of DNA strands.

Amplified oligos were sequenced on a commercial Illumina Hiseq 4000 platform with 150 paired-end cycles (Section S5) and then deeply analyzed by a set of statistics methods developed from bioinformatic BLAST program (fig. S3 and Section S6). Sequencing analysis revealed various errors, including base substitution and indel. Among them, single letter error accounted for the majority in mutant sequenced reads for both PCR (80.1%) and iDR (81.7%) (Fig. 2E and fig. S4). The total indel (0.03%) and substitution (0.2%) rate were consistent with previous studies(*13*). RS code embedded in BASIC code is able to correct multiple errors in the same DNA strand, but will also largely increase the computation complexity (*17, 26*). Therefore, both sequenced reads with no or single substitution/indel errors were collected as valid reads for further analysis and decoding (fig. S1B).

The distribution of the number of reads per each given sequence were different between iDR and PCR of various thermal cycles of from 10 to 60. The distribution normality was quantified by a modified mathematical statistics function (Section S6). In theory, oligo pool of higher value distribution normality requires less sequencing reads to recover all sequences (*12*). The coverage distribution of both PCR and iDR were positively skewed with a long tail, which comprising of high copy number oligos, the normality of PCR decreased from 0.50 to 0.28 with thermal cycles increased from 10 to 60 (Fig. 2F). In comparison, a normality of 0.60 was obtained for iDR, which indicated a more balanced amplification of DNA pool. The proportion of oligos with high copy numbers (i.e. the top 30% high coverage oligos) continued growing with more PCR thermal cycles (Fig. 2G). Additionally, the dropout rate increased significantly from 4.18% to 13.80% with PCR amplification becoming deeper, much higher than the 1.33% dropout rate of iDR. No obvious difference was observed between the coverage distribution of iDR amplification from free oligo pool and oligo pool immobilized on magnetic beads (fig. S5).

In the steps of sequencing and information decoding, the DNA sequences with more copy numbers are considered to be likely correct with higher probability. Therefore, deep errors, reads with mutants in high copy numbers, accumulated in biased amplification and impeded information decoding. Sequencing reads were further sorted out as group of M0G0 (with no letter error) and M1G1 (with single letter substitution or indel error) by developed BLAST programs. The two groups were mapped to the original designed oligo sequence and the copy number for every single reference oligo was counted respectively. For iDR, average 2 oligos were identified with copy number in M1G1 higher than M0G0. This oligo could be considered as accumulated deep error, but average 15, 29 and 79 oligos were identified in 10c, 30c and 60c sample respectively (Fig. 2H). It clearly demonstrated that the error was prone to accumulate in biased amplification process.

Low replication fidelity of DNA polymerases and around 1% miss reading in NGS sequencing process(*27*) largely contributed to the massive error reads associated with low-abundant DNA strands, e.g. 1-2 copy numbers, but it is relatively easy to identify them from correct reads in high copy number. In PCR amplification, product-as-template and inefficient priming process caused the high amplification bias and made PCR amplification prone to error-spreading and oligo dropout.

In contrast, iDR was designed to synthesize new oligo only from the original templates without priming process for replication initiation and therefore iDR achieved a low biased and error-spreading proof amplification and will require much less calculation resource in decoding process. Additionally, high temperature treatment resulted to DNA oligo decay especially for long strands(*16, 17*). The decay lost rate in a 10 cycles PCR was calculated as 21.8‰ and 0.035‰ for iDR (Fig. 2B and Section S7). Finally, all the files in DNA pool was successfully decoded following the BASIC decoding process and the minimal mean coverage depth of sequencing reads were calculated (Table S2).

### Sustainable DNA storage for repeated reading

Repeatable reading is crucial for practical DNA storage system. So far, different repeated reading systems have been proposed and tested, in which generally DNA pool was amplified from aliquot of previous amplified product (Fig. 3A). However, error rate and oligo dropout were significantly accumulated through this repeated amplification(*12, 13, 16*). Due to its low biased amplification, we believe that the iDR system is ideal for robust repeated reading in DNA storage. In this isothermal system (Fig. 3B), we propose to use both the low biased iDR amplification and DNA pool immobilization on microbeads to achieve stable and repeatable reading.

**Fig. 3.**
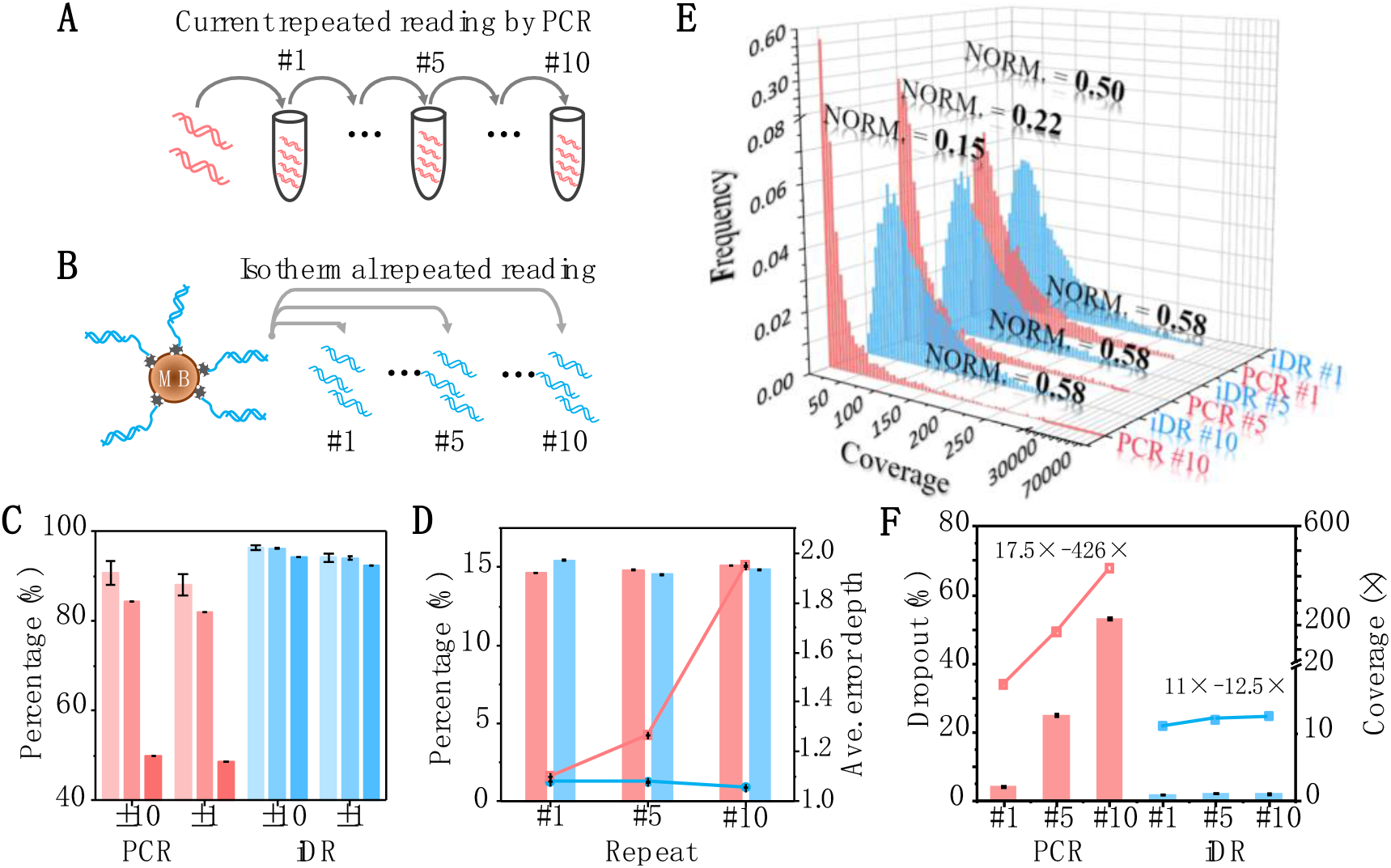
Multiple information reading from DNA storage. **(A)** Illustration of successive DNA reading. Repeated PCR amplification was successively performed 10 times with aliquot of previous reaction as template. **(B)** the oligos pool immobilized on magnetic beads was successive amplified 10 times by iDR. The same master oligo pool was used as the initial template for PCR and iDR and #1, #5 and #10 amplified oligos were analyzed. **(C)** The proportion of reads with up to 10 substitution or indel letter error per million sequenced reads were counted as 90.65% (#1 PCR, light red), 84.25% (#5 PCR, red), 49.97% (#10 PCR, dark red), 96.35% (#1 iDR, light blue), 96.14% (#5 iDR, blue), and 94.12% (#10 iDR, dark blue), and error reads with up to 1 error were counted as 88.18% (#1 PCR), 82.05% (#5 PCR), 48.58% (#1 PCR), 94.18% (#1 iDR), 94.04% (#5 iDR), and 92.24% (#10 iDR). Error bars represent the mean ±s.d., where n = 3. **(D)** The proportion of M1G1 reads per million valid reads were, PCR (red column), 14.65% (#1 PCR), 14.82% (#5 PCR), 15.10% (#10 PCR), and iDR (blue column) 15.46% (#1 iDR), 14.55% (#5 iDR), and 14.86% (#10 iDR). The average error reads depth of PCR (red line, 1.10 for #1, 1.26 for #5, 1.95 for #10) increased and iDR (blue), but iDR (blue line, 1.08 for #1, 1.08 for #5, 1.05 for #10) remained stable. Error bars represent the mean ±s.d., where n = 3. **(E)** The distribution of reads number per each given sequence per million sequenced reads of #1, #5, and #10 of PCR and iDR. The distribution normality was 0.5 (#1 PCR), 0.22 (#5 PCR), 0.15 (#10 PCR), 0.58 (#1, #5, #10 iDR) respectively. **(F)** The dropout rate for random sequenced reads set with 10x coverage depth was plotted, PCR (red column, 4.18% for #1, 22.89% for #5 and 53.19% for #10) and iDR (blue column, 1.86% for #1, 2.08% for #5 and 2.09% for #10). Error bars represent the mean ±s.d., where n = 3. The coverage depth for random sequenced reads with 1.56% dropout was calculated as 17.2 (#1 PCR), 167 (#5 PCR), 426 (#10 PCR) and 11 (#1 iDR), 12 (#5 iDR), and 12.5 (#10 iDR) respectively. Error bars represent the mean ±s.d., where n = 10.

The amplification capability for repeated deep reading oligo pool containing from 11,520 to 89,088 DNA strands was examined. PCR amplification was successively performed 10 times from aliquot of previous reaction and iDR amplification was repeated 10 times from immobilized oligos pool. In #1, #5 and #10 of successive PCR, the proportion of amplified oligos with up to total 10 substitution or indel letter error decreased from 90.65% (±10 of PCR #1) to 49.97% (±10 of PCR #10) and from 88.18% (±1 of PCR #1) to 48.58% (±1 of PCR #10). In comparison, the repeated iDR remained very consistent over 90% (Fig. 3C). These statistics results indicated that large noise was introduced during PCR procedure and amplified oligo with imperfect length increased from inefficient replication or miss priming. No obvious difference was observed in proportion of M1G1 in total sequenced reads between PCR and iDR, except that the mean copy number in M1G1 reads increased from 1.10 of #1 PCR to 1.95 of #10 PCR, but only slightly changed for iDR, 1.08 of #1, 1.08 of #5 and 1.05 of #10 (Fig. 3D). This result was in agreement with previous experiment and indicated that error accumulated to high copy number during PCR and iDR achieved error-spreading proof amplification, in which amplified DNA strand with error was not further amplified as template.

Interestingly, it was observed that the distribution of the number of reads per each given sequence changed differently. Normality of #1, #5 and #10 PCR decreased significantly from 0.50 to 0.15. In contrast, #1, #5 and #10 iDR gave a consistent normality of 0.58 (Fig. 3E). For successive PCR, the coverage distribution was largely positively skewed and the proportion of both low copy number and high copy number oligos significantly increased. It indicated that large enrichment driven by amplification bias for part of oligos efficiently occurred with the successive PCR, but not in repeated iDR. Furthermore, it was observed that only top 1% oligo of high coverage increased its proportion significantly and both 1% oligos of middle and low coverage decreased while the oligo pool was successively read by PCR. In contrast, all the 1% oligos remained steady in repeated iDR (fig. S6). The dropout rate in random valid reads set with 10x coverage was quantified to further assess the amplification bias. For PCR, the dropout rate increased sharply from 4.18% (#1 PCR) to 53.19% (#10 PCR) (Fig. 3F), but iDR remained steady at about 2%. Because the BASIC encoding algorithm has a 1.56% dropout rate, we also calculated the coverage depth for random valid reads with 1.56% dropout rate (Section S3), the crucial parameter for the theoretical minimum decoding coverage. For PCR, the minimum coverage depth was quantified as 17.2x (#1 PCR) and 167x (#5 PCR) and 426x (#10 PCR). Here, the minimum decoding coverage (426x) for 10# PCR was calculated by fitting curve of the coverage-dropout ratio (Section S8). For iDR, the minimum coverage depth was quantified as 11 for #1, 12 for #5 and 12.5 for #10. Therefore, for 445KB file encoded in 11,520 DNA oligos, 10,101,935 and 156,115 minimal sequencing reads were necessary for perfect decoding (Fig. 4 and Section S8).

**Fig. 4.**
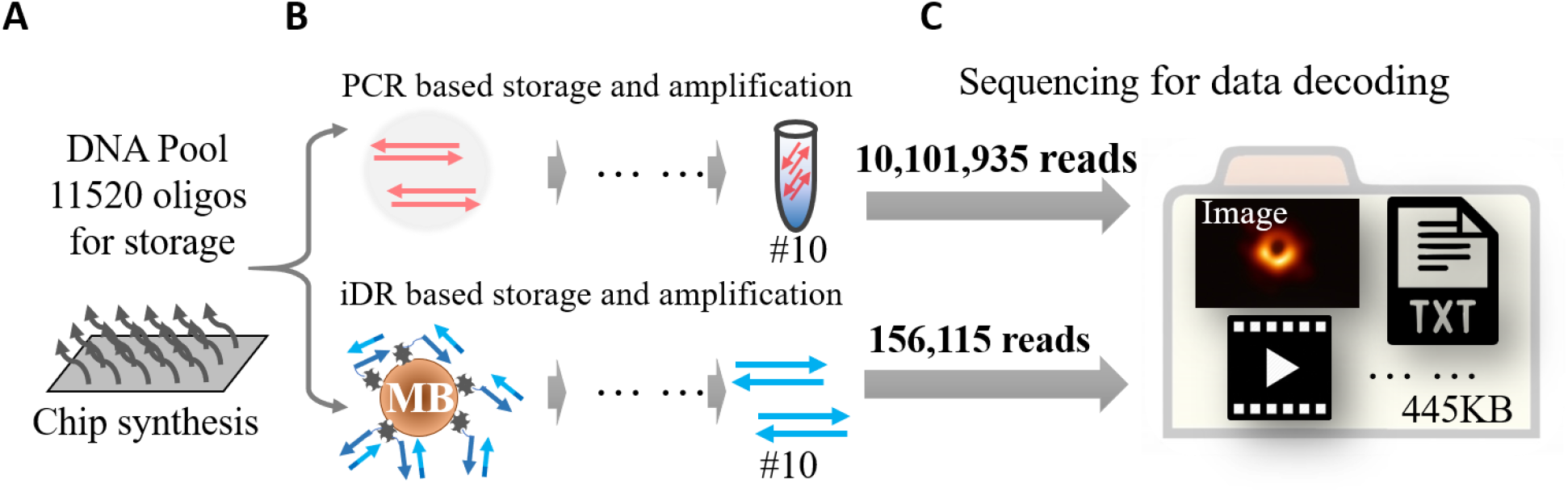
Sustainable and robust iDR DNA storage. **(A)** Chip-synthesized DNA oligo pool, in which stores digital information including image, video, and text. **(B)** The DNA pool was repeatedly amplified PCR or iDR, respectively. **(C)** For current reported DNA system with repeated reading based on PCR, 10,101,935 NGS noise reads was necessary for perfect decoding 445KB digital files, but only 156,115 NGS noise reads were necessary for iDR.

Thus far, for PCR based sustainable DNA media reading, one strategy is deep amplification from trace DNA material with large number of thermal cycles and another is successive amplification from aliquot of previous reaction. However, we demonstrated that both strategies are not practical. The nature of PCR including especially inefficient priming and product-as-template generated large amplification bias that significantly skewed the copy number distribution and resulted huge dropout in both 60 cycles deep amplification and 10 times successive amplification, especially for oligo pool with huge sequence complexity. Based on these deep statistical analyses, we pointed out that iDR amplification of immobilized DNA pool was stable, robust and suitable for repeated reading in DNA data storage.

## Discussion

The concept of DNA data storage has been proposed for a long time(*28*). However, practical DNA data storage requires more powerful capability for all related biotechnologies, including synthesis, sequencing and manipulation of oligo pool with high sequence complexity. Not until lately, the significantly improved capability in DNA synthesis and sequencing started making it a distinct reality. In many respects, the size of oligo pool used in DNA storage is already several orders of magnitude larger than that in other applications. However, there is still a lack of technology specifically tailored for DNA storage, especially for the manipulation of DNA oligo pool with huge sequence complexity and unevenness of molecule copy numbers. In studies to date, PCR is still the only method for amplification of oligo pool for reading information. However, with deep bioinformatic and statistics analysis, we demonstrated that PCR amplification skewed the oligo pool even after a very few thermal cycles (10 cycles) and the skewness largely increased, deep error spreading and massive dropout, as the function of amplification cycle numbers. Deep error, mutant sequenced reads in high coverage, interfered decoding and caused significant increased calculation, but decoding fatally crashed due to massive dropout. We contend that the features of low temperature, priming-free, and enzyme-mediated double helix unzipping in iDR amplification presented here, overcame the major amplification bias related issues of PCR including the inefficient priming, product-as-template, sequence context dependent, and high temperature heating. iDR achieved very stable amplification performance, which efficiently prevented the mutant error from spreading, decreased over 70% deep error, and achieved successive repeated deep reading with consistent outcome quality. Actually, the physical storage density, the most significant advantage of DNA storage with a calculation of a few kilograms DNA material for storage of data from the whole human society(*29*), highly depend on the quality of DNA reading. Therefore, higher physical storage density could be achieved by iDR system, which could store information with at least two orders of magnitude less DNA material and sequencing resource than the current PCR method (Fig. 4 and Section S8).

Although there are many variants of PCR, such as emulsion PCR, digital PCR, and multiplex PCR, none of them can avoid the crucial issues of product-as-template, inefficient priming, and complex thermal regulation. Each of these will raised the cost for handling the large DNA pool. Our iDR is adapted from strand displacement replication, but this is the first system applying this amplification mechanism on DNA pool with high sequence complexity for artificial information storage and demonstrated that immobilized DNA pool read by iDR is able to stably read data deeply and repeatedly. Originally, the core thing of DNA storage is adapting current biotechnology for artificial purpose of storing information at a large scale and accuracy which is distinct for bioengineering approaches. Undeniably, PCR possesses power in DNA amplification, and iDR could work with PCR together in the different steps of DNA storage, e.g., material preparation and stably reading respectively.

Furthermore, Replication fidelity of DNAP I Klenow fragment used in iDR system is much lower than Q5 DNA polymerase in PCR. There is still much room for improvement. Additionally, practical DNA storage system must move out of biochemical test tube to build device by highly integrating all related biochemical processes. Prototypes of DNA storage hardware have already been proposed with high density DNA material in microfluidic device(*30, 31*). We contend that besides some crucial features of iDR make it more fit for hardware construction. In iDR, only low temperature, slightly higher than room temperature, was required. Considering the operation of DNA storage up to large scale, there will be a huge difference in energy consumption. Furthermore, iDR is able to work in a primer-free fashion, only defined protein enzyme mix is required no matter what information was encoded, which makes it possible for universal information reading without any sequence information required in advance. To the best of our knowledge, iDR is the first system for stable repeated DNA information reading which is crucial and necessary feature for practical and sustainable storage hardware. Therefore, we believed that this new biochemical framework with stable repeated reading lays a foundation for development of practical and sustainable DNA storage.

## Materials and Methods

### Preparation of DNA Master Oligo Pool

The pool (Pool 1/2--Twist Bioscience; Pool 3--CustomArray) was resuspended in 1x TE buffer for a final concentration of 2 ng/uL. PCR was performed using Q5 High-Fidelity DNA Polymerase (New England Biolabs, NEB). Mix 10 ng of ssDNA pool (5 uL) with 2 μL of 100 μM of the Adaptor I and 2 μL of 100 μM of the Adaptor II (Adaptor II-1), 10 μL 5x Q5 Reaction Buffer, 4 μL of 2.5 mM dNTPs, 0.5 μL Q5 High-Fidelity DNA Polymerase. Thermocycling conditions were as follows: 5 min at 98°C; 10 cycles of: 30 s at 98°C, 30 s at 58°C, 15 s at 72°C; followed by a 5 min extension at 72°C. The reaction was then purified according to the instructions in the Eastep Gel and PCR Cleanup Kit and eluted in 50 μL DNase/RNase-free water. This library was considered the master pool. All primers we used were synthesized by GENEWIZ, which are listed in table S3.

### iDR reaction

10 ng of DNA oligo out of the master pool was attached to 1 μL of Streptavidin Magnetic Beads (NEB). The iDR reaction mixtures contained 1 μL of the DNA template attached to the beads (10 ng/μL), 0.25 mM dNTPs, 2.5 μL 10x NEBuffer 2, 0.08 U/μL Nt.BbvCI (NEB), 0.16 U/μL KF polymerase (exo-) (Vazyme), 4 μM T4 Gene 32 Protein (gp32), 0.2 mg/mL BSA, 0.5 μM Adaptor II (For production of ssDNA, adaptor II was not added.). The mixtures were incubated at 37°C for 30 min. After amplification, the specific amplicons and the templates were isolated through magnetic pull-down. In the process of repeated iDR, the templates attached to magnetic beads were washed by Wash/Binding buffer (0.5 M NaCl, 20 mM Tris-HCl (pH 7.5), 1 mM EDTA) twice and mixed with the above mentioned iDR reaction mixtures except for the DNA templates. The processes were proceeded for 10 times. To retrieve the information, the amplified products were purified and then sequenced on one Illumina Hiseq 4000 platform with 150 paired-end cycles in Novogene (Section S5).

### PCR reaction

PCR was performed using Q5 High-Fidelity DNA Polymerase and adaptor I (pool 2/3) or adaptor I-1 (pool 1)/adaptor II (10 ng DNA master pool, 2 μL of adaptor I(100 μM); 2 μL of adaptor II (100 μM)), 10 μL 5x Q5 reaction buffer in a 50 μL reaction. Thermocycling conditions were as follows: 5 min at 98°C; 10 cycles of: 30 s at 98°C, 30 s at 58°C, 10 s at 72°C, followed by extension at 72°C for 5 min. In the repeated PCR, each subsequent PCR reaction consumed 1 μL of the prior PCR reaction and employed 10 cycles in each 50 μL reaction. The PCR product was purified by Eastep Gel and PCR Cleanup Kit and eluted in 50 μL DNase/RNase-free water. Then we sequenced the PCR product on Illumina Hiseq 4000 platform. For deep PCR, the reaction conditions were the same as above mentioned except cycles number.

## Acknowledgments

We would like to thank Professor Hanxu Hou from Dongguan University of Technology for advice and assistance with designing algorithm for BASIC code. We also thank Yixi Wang from College of intelligence and computing at Tianjin University for her help in test of encoding system.

## Author contributions

Y.G. and H.Q. designed and performed all experiments. X.C., J.H. and C.Z. designed and developed the encoding and decoding program. X.C. developed program for the bioinformatics statistics analysis. Y.G., H.Qiao and H.Q. collected and analyzed all experiment data. Y.K. participated in the discussion of experiment data. Y.G., Y.K. and H.Q. wrote the manuscript. H.Q. designed experiments, analyzed data and supervised this work.

## Data and materials availability

The BASIC code for encoding and decoding for both Linux and Windows and bioinformatic analysis programs may be obtained via (https://github.com/xiaomingao/DNA-information-storage). Furthermore, the original sequencing FASTQ file and the designed sequence file may be obtained via (http://pan.tju.edu.cn:80/#/link/627DB5C9EB819984F1183D8D4A0B72E3, Code: oZ3D).

## References

1. S. Kosuri, G. M. Church, Large-scale de novo DNA synthesis: technologies and applications. Nat Methods 11, 499–507 (2014).

2. S. Goodwin, J. D. McPherson, W. R. McCombie, Coming of age: ten years of next-generation sequencing technologies. Nat Rev Genet 17, 333–351 (2016).

3. I. Kozarewa, Z. Ning, M. A. Quail, M. J. Sanders, M. Berriman, D. J. Turner, Amplification-free Illumina sequencing-library preparation facilitates improved mapping and assembly of (G+C)-biased genomes. Nat Methods 6, 291–295 (2009).

4. L. Organick, Y. J. Chen, S. Dumas Ang, R. Lopez, X. Liu, K. Strauss, L. Ceze, Probing the physical limits of reliable DNA data retrieval. Nat Commun 11, 616 (2020).

5. Y.-J. Chen, C. N. Takahashi, L. Organick, K. Stewart, S. D. Ang, P. Weiss, B. Peck, G. Seelig, L. Ceze, K. Strauss, Quantifying Molecular Bias in DNA Data Storage. (2019).

6. J. C. Klein, M. J. Lajoie, J. J. Schwartz, E. M. Strauch, J. Nelson, D. Baker, J. Shendure, Multiplex pairwise assembly of array-derived DNA oligonucleotides. Nucleic Acids Res 44, e43 (2016).

7. H. G. Jingdong Tian, Nijing Sheng, Xiaochuan Zhou, Erdogan Gulari, Xiaolian Gao & George Church, Accurate multiplex gene synthesis from programmable DNA microchips. Nature 432, 1050–1054 (2004).

8. S. Linnarsson, Recent advances in DNA sequencing methods - general principles of sample preparation. Exp Cell Res 316, 1339–1343 (2010).

9. S. Kosuri, N. Eroshenko, E. M. Leproust, M. Super, J. Way, J. B. Li, G. M. Church, Scalable gene synthesis by selective amplification of DNA pools from high-fidelity microchips. Nat Biotechnol 28, 1295–1299 (2010).

10. R. J. Lipshutz, S. P. Fodor, T. R. Gingeras, D. J. Lockhart, High density synthetic oligonucleotide arrays. Nat Genet 21, 20–24 (1999).

11. M. T. Bonde, S. Kosuri, H. J. Genee, K. Sarup-Lytzen, G. M. Church, M. O. Sommer, H. H. Wang, Direct mutagenesis of thousands of genomic targets using microarray-derived oligonucleotides. ACS Synth Biol 4, 17–22 (2015).

12. Y. Erlich, D. Zielinski, DNA Fountain enables a robust and efficient storage architecture. Science 355, 950–954 (2017).

13. L. Organick, S. D. Ang, Y. J. Chen, R. Lopez, S. Yekhanin, K. Makarychev, M. Z. Racz, G. Kamath, P. Gopalan, B. Nguyen, C. N. Takahashi, S. Newman, H. Y. Parker, C. Rashtchian, K. Stewart, G. Gupta, R. Carlson, J. Mulligan, D. Carmean, G. Seelig, L. Ceze, K. Strauss, Random access in large-scale DNA data storage. Nat Biotechnol 36, 242–248 (2018).

14. D. Aird, M. G. Ross, W. S. Chen, M. Danielsson, T. Fennell, C. Russ, D. B. Jaffe, C. Nusbaum, A. Gnirke, Analyzing and minimizing PCR amplification bias in Illumina sequencing libraries. Genome Biol 12, R18 (2011).

15. N. Goldman, P. Bertone, S. Chen, C. Dessimoz, E. M. LeProust, B. Sipos, E. Birney, Towards practical, high-capacity, low-maintenance information storage in synthesized DNA. Nature 494, 77–80 (2013).

16. R. Heckel, G. Mikutis, R. N. Grass, A Characterization of the DNA Data Storage Channel. Sci Rep 9, 9663 (2019).

17. R. N. Grass, R. Heckel, M. Puddu, D. Paunescu, W. J. Stark, Robust chemical preservation of digital information on DNA in silica with error-correcting codes. Angewandte Chemie 54, 2552–2555 (2015).

18. H. Hou, K. W. Shum, M. Chen, H. Li, BASIC Codes: Low-Complexity Regenerating Codes for Distributed Storage Systems. IEEE Transactions on Information Theory 62, 3053–3069 (2016).

19. H. Hou, Y. S. Han, K. W. Shum, H. Li, A Unified Form of EVENODD and RDP Codes and Their Efficient Decoding. IEEE Transactions on Communications 66, 5053–5066 (2018).

20. L. Organick, Y.-J. Chen, S. D. Ang, R. Lopez, K. Strauss, L. Ceze, Experimental Assessment of PCR Specificity and Copy Number for Reliable Data Retrieval in DNA Storage. (2019).

21. E. L. van Dijk, Y. Jaszczyszyn, C. Thermes, Library preparation methods for next-generation sequencing: tone down the bias. Exp Cell Res 322, 12–20 (2014).

22. J. Dabney, M. Meyer, Length and GC-biases during sequencing library amplification: a comparison of various polymerase-buffer systems with ancient and modern DNA sequencing libraries. Biotechniques 52, 87–94 (2012).

23. Y. Zhao, F. Chen, Q. Li, L. Wang, C. Fan, Isothermal Amplification of Nucleic Acids. Chem Rev 115, 12491–12545 (2015).

24. M. S. Reid, X. C. Le, H. Zhang, Exponential Isothermal Amplification of Nucleic Acids and Assays for Proteins, Cells, Small Molecules, and Enzyme Activities: An EXPAR Example. Angewandte Chemie 57, 11856–11866 (2018).

25. D. Y. Zhang, G. Seelig, Dynamic DNA nanotechnology using strand-displacement reactions. Nat Chem 3, 103–113 (2011).

26. L. Anavy, I. Vaknin, O. Atar, R. Amit, Z. Yakhini, Data storage in DNA with fewer synthesis cycles using composite DNA letters. Nat Biotechnol, (2019).

27. S. M. Yazdi, Y. Yuan, J. Ma, H. Zhao, O. Milenkovic, A Rewritable, Random-Access DNA-Based Storage System. Sci Rep 5, 14138 (2015).

28. M. R. Wallace, Molecular Cybernetics: The Next Step? Kybernetes 7, 265–268 (1978).

29. A. Extance, How DNA could store all the world’s data. Nature 537, 22–24 (2016).

30. S. Newman, A. P. Stephenson, M. Willsey, B. H. Nguyen, C. N. Takahashi, K. Strauss, L. Ceze, High density DNA data storage library via dehydration with digital microfluidic retrieval. Nat Commun 10, 1706 (2019).

31. C. N. Takahashi, B. H. Nguyen, K. Strauss, L. Ceze, Demonstration of End-to-End Automation of DNA Data Storage. Sci Rep 9, 4998 (2019).

