## Supplementary for "Low-Bias Amplification for Robust DNA Data Readout"

### Section S1. BASIC code strategies.

**Distributed Storage Systems in application to DNA storage.** DNA coding is a new type of distributed storage system. In a distributed storage system, data redundancy is the most basic strategy for ensuring system reliability and improving data availability. By storing multiple instances of the same data file with different nodes to ensure data availability, even some of the data is unavailable, the remaining nodes can still reconstruct the original data. Redundancy strategies must consider two points: firstly, how to create redundant data, and secondly, how to reconstruct data when some nodes fail. Currently, widely used redundancy strategies are replication and erasure coding. Replication distributes multiple copies of a file to different nodes in the system. As long as one of these copies is valid, the whole file can be obtained. This method has high reading and writing efficiency, but it has low storage utilization and is not suitable for DNA coding. Erasure Coding is another important redundancy strategy. (N, K) erasure code matrix divides an original file of size M into K blocks, each block with size of M/K; then the K block files are encoded into N code blocks and distributed to N nodes. The original file can be reconstructed from any K code blocks in the N code blocks. Erasure coding requires less storage than replication, whereas the calculation is relatively complex. BASIC code is a kind of distributed erasure code designed for DNA coding, aiming at maximizing storage utilization and effectively guaranteeing the reliability of the storage system.

This section briefly introduces the binary cyclic code. Let p be a prime number greater than N and the ring $R_p$ is defined as $R_p \coloneqq \mathbb{F}_2[z]/(1+2^p)$, where the element $\sum_{i=0}^{p-1} a_i z^i$ in $R_p$ is called polynomial, and the vector $(a_0, a_1, \dots, a_{p-1}) \in \mathbb{F}_2^p$ is a codeword of polynomial $\sum_{i=0}^{p-1} a_i z^i$. A binary cyclic code of length p is a subset of additions and z-multiplier closures defined in $R_p$, where addition is an XOR operation and z multiplication is a cyclic right shift operation.

In this paper, we consider the parity code $\mathcal{C}_p$, which represents the set of polynomials for which all non-zero coefficient entries in $R_p$ are even. $\mathcal{C}_p$ is formalized as:

$$\mathcal{C}_p \coloneqq \{\sum_{i=0}^{p-1} a_i z^i \in R_p | a_0 + a_1 + \cdots + a_{p-1} = 0\}$$

$$= \{a(z) \in R_p | a(z) \equiv 0 \; mod \; (1+z)\}$$

$$= \{a(z)(1+z) \big| a(z) \in R_p\}$$

The coefficient of the highest term of the element a(z) in $\mathcal{C}_p$ is the sum of the former m-1 coefficients, i.e., $a_{p-1} = \sum_{i=0}^{p-2} a_i$ . It can be verified that $\mathcal{C}_p$ satisfies the addition and z multiplication closure. Because the operations in the domain $\mathcal{C}_p$ are only XOR and the right-shifted loop, it can be well applied to design the encoding system.

The addition defined in $\mathcal{C}_p$ is an XOR operation, which will not be described in detail here. The z-multiplication operation in $\mathcal{C}_p$ is defined as $\mathrm{z}_p: R_p \times C_p \to C_p$. $\mathrm{z}_p(2^i, \mathrm{c}) = 2^i * c \; mod(2^p + 1)$, that is, the loop shifts right by i bits. For example, the codeword is $\mathrm{z}_3(2^1, 101) = 011$; $\mathrm{z}_3(2^1 + 2^0, 101) = \mathrm{z}_3(2^1, 101) + \mathrm{z}_3(2^0, 101) = 011 + 101 = 110$.

Vandermonde matrix is $\mathrm{V} = [\mathrm{v}_{i,j}]_{n \times k}$, where $\mathrm{v}_{i,j} = \alpha_j^{i-1}$. The determinant of an N-order Vandermonde square matrix can be represented as $\det(\mathrm{V}) = \prod_{1 \le i < j \le n} (\alpha_j - \alpha_i)$. When $\alpha_i$ are not the same, $\det(\mathrm{V})$ is not zero. The Vandermonde matrix has many interesting features. The most important thing here is that the sub-polynomials formed by any row and column are invertible. The Vandermonde matrix and its transformation matrix ensure that the encoding data can be decoded. For convenience, note that $\mathrm{v}_i = (\mathrm{v}_{i,1}, \mathrm{v}_{i,2}, \dots, \mathrm{v}_{i,n}) = (\alpha_1^{i-1}, \alpha_2^{i-1}, \dots, \alpha_n^{i-1})$ is the ith row vector of V.

**Constraints in encoding DNA digital information.** GC content of DNA sequence in payload was restricted in the range between 30% and 70%. In fact, GC content of whole sequence including payload and adaptor was in the range between 35% and 66% in our all decoding sequences. Meanwhile, long homopolymers (i.e., AAAA, TTTT, GGGG, CCCC) were dropped. More than 6-bp self-sequence complementarity and 10bp inter-sequence complementarity were avoided. Certain sequences were circumvented (table S4).

**The distributed storage system coding strategy.** The goal was to transform the input file to DNA sequence reads with biochemical constraints. DNA basic code should enable error-detection, error-correction and full recovery. There were two key steps: (a) erasure coding, (b) RS coding. Since the sequence reads needed to satisfy the biochemical constraints, both processes included the step of filtering the sequences.

**Erasure coding.** The algorithm divided the file into non-overlapping groups of length K*L bits, each group containing K binary data of length L. In the subsequent encoding step, the algorithm processed the data in groups. K and L were arbitrarily defined. Here we used K=252, L=256 bits (32 bytes) because this parameter setting was compatible with the standard computing environment and was within the capabilities of actual manufacturing. The BASIC encoding was shown as follows.

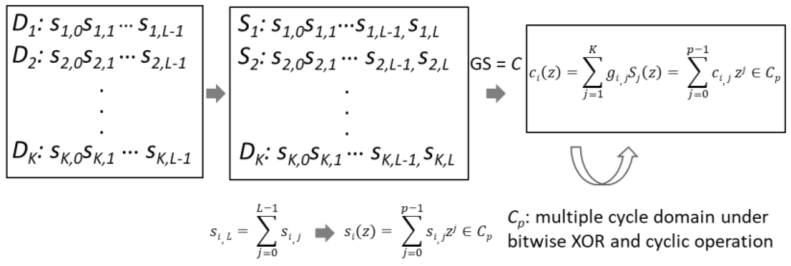


For each group, we proposed BASIC code to encode the K pieces of data one by one. The BASIC code used codeword as the encoding and decoding unit, a codeword was a binary data of L bits. K codewords of length L were encoded into N codewords by polynomial matrix operations on the polynomial cycle domain $\mathcal{C}_p$, where N>K, N-K was the maximum number of error sequences per group. Actually, N was 256, which meant each group of data allowed a maximum of 4 reads to be lost or corrupted. For simplicity of discussion, a codewordis represented as $D_i$ or $C_i$. The input data was regarded as a vector $\mathrm{D} = (\mathrm{D}_1, \mathrm{D}_2, \ldots, \mathrm{D}_K)$, and the encoded data was a vector $\mathrm{C} = (\mathrm{C}_1, C_2, \ldots, C_N)$. The generation matrix G is a matrix defined on $R_p$, which was an improvement of the Vandermonde matrix. The reason for using the Vandermonde matrix was that the BASIC data recovery algorithm required the encoding matrix to be reversible by any n*n submatrix. The specific process was as follows:

1) In order to ensure that the obtained $s_i(z)$ was in $C_p$, for a set of K*L data $\mathrm{D} = (\mathrm{D}_1, \mathrm{D}_2, \ldots, \mathrm{D}_K)$, first added a parity bit at the end of each sequence read. The parity bit was calculated as $s_{i,p-1} = \sum_{j=0}^{p-2} s_{i,j}$ (where the addition is an exclusive-OR operation). After that, the codeword was S= $(S_1, S_2, \ldots, S_K)$. And p was 257 here.

2) Initialize the generation matrix G as follows:

$$\mathrm{G} = \begin{bmatrix} 2^0 & 2^1 & \cdots & 2^{K-1} \\ 2^{0\times 2} & 2^{1\times 2} & & 2^{(K-1)\times 2} \\ \vdots & & \ddots & \vdots \\ 2^{0\times N} & 2^{1\times N} & \cdots & 2^{(K-1)\times N} \end{bmatrix} mod\ (2^p + 1) = [g_{i,j} = 2^{(j-1)\times i\ mod\ p}]_{N\times K}$$

First, a constant S was added to all codewords, because zero-by-zero multiplication was still zero, which could cause the algorithm to not terminate. Next, initialized $C_i$, got $C_i \leftarrow g_i S^T$,$g_i = (g_{i,1}, g_{i,2}, \ldots, g_{i,K})$ was the i-th row of G. Finally, for any $C_i$, the last parity bit was removed and its first L bits were saved.

**RS Encoding.** In order to ensure the accuracy of data storage in DNA, RS codes were used to increase error correction and repair capabilities. Here, for each sequence reads, 2 bytes were allocated for the RS codes, which could detect errors within 2 bytes and correct errors within 1 byte.

**Filtering.** Each codeword C*i* consisted of three parts: the sequence number of the group, the sequence number within the group, and the number of adjustments to $\mathrm{G}_i$. The index part was allocated a total of 4 bytes, in which the first 8 bits represented the group address, the middle 8 bits represented the address within each group, and the last 16 bits represented the number of adjustments of $\mathrm{G}_i$.

All sequence reads must meet the biochemical constraints, which meant that the sequence could not contain any avoidance sequences. For the sequences obtained after the erasure coding, we generated a number of random equal length sequences, so that the encoding sequences didn't contain any avoidance sequence after exclusive-OR operation with one of the random sequences. It could be proved by experiments that the upper 8 bits of the 16 bits, which represent the number of adjustments of $\mathrm{G}_i$, were always 0. Thus, we used these bits to store the sequence number of the random sequence that was XORed for each encoding sequence. When decoding, it was only necessary to find a random sequence according to the serial number for XOR. The same process was performed for the individual RS codes. A number of random sequences with the same length of the RS code were generated for XOR operation. Since there were no bits to store the sequence number of the RS XOR sequence, we built a mapping table to store the RS code and its corresponding sequence number. It should be noted that all RS codes in the mapping table were unique.

**Decoding.** The decoding process was reversed step by step according to the encoding process. XOR operation was performed according to the mapping table to restore the RS code, and then the RS code was used for error correction to ensure that each sequence was accurate. Restore the BASIC code sequence. For each group of data, it was decoded according to the BASIC decoding algorithm.

1) Find K lossless sequences for each group. A parity bit is added for each codeword, which is denoted as $\mathrm{C}' = (\mathrm{C}'_1, \mathrm{C}'_2, \ldots, \mathrm{C}'_K)$.

2) Construct a generation matrix G' according to the intragroup address and the number of adjustments of $G_i$. Calculate G's inverse matrix $G'^{-1} = [f_{i,j}]_{K\times K}$.

3) Decode $D'_i = \sum_{j=1}^{k} f_{i,j} * C'_j$ according to the matrix $G'^{-1}$. Remove the last parity bit and finally recover the original data.

The decoding processes are shown as follows.

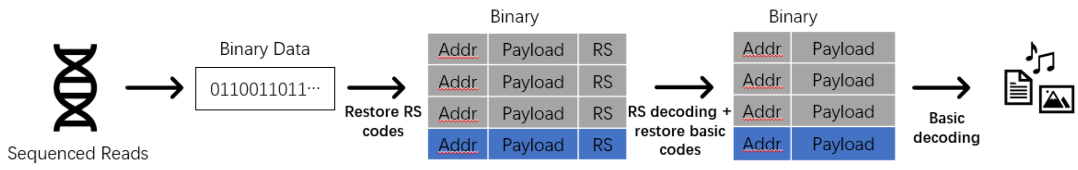

**Section S2. Genome sequence were converted to binary information.**

Human mitochondrial genome containing Chinese (16,570 bp), Italy (16,569 bp), Native American (16,570 bp), and South African (16,567 bp) were converted to binary data by a simple rule of T to 00, G to 01, C to 10 and A to 11. Genome information comes from mtDB – Human Mitochondrial Genome Database (http://www.mtdb.igp.uu.se/sequences.php). And then the binary file was encoded to oligo pool1/2 (Twist Bioscience).

The 1.08-Mbp *M. mycoides* JCVI-syn1.0 genome sequence (accession CP002027) were also converted and encoded to oligo pool 3 (CustomArray).

### Section S3. Calculating logical density and redundancy for DNA storage systems.

Logical density calculated for previous studies is taken from Organick et al.. In this work, due to adjustable system parameter K and L, we take standard system parameter (K=252, L=256) as example. Here, we have used 10,752 DNA sequences (a part of DNA sequences in Pool 2) of length 200 with payload of length 152 to store 329KB of the compressed file yielding a net density of approximately 1.65 bits per base (1.25 bits per base including primers). According to our encoding strategy which allows a maximum of 4 DNA sequences to be lost or corrupted in each group (each group contains 256 DNA sequences), the redundancy can be calculated ideally as: 4/256=1.56%.

**Section S4. Energy consumption.**

We assumed that the system was thermally insulated and there was no energy consumption during the state of constant temperature. Energy consumption Q = c * v * ($T_2$-$T_1$) (c is the specific heat capacity of the solution; the volume of the system is v; $T_1$, $T_2$ is the initial and the final temperature respectively) when the reaction is exothermic. Energy consumption Q = c * v * ($T_1$-$T_2$) while the reaction is endothermic.

We took 25 μL of water as an example. We supposed that the specific heat capacity is equivalent to water's (c, c = 4.2 * $10^3$ J/(kg * ℃)) and the initial temperature ($T_0$) is 25℃. The thermocycling conditions of PCR were as follows: 5 min at 98℃; 10 cycles of: 30s at 98℃, 30s at 58℃, 10s at 72℃, followed by a 5 min.

25-98 Q = 4.2 * $10^3$ * 25 *$10^{-6}$ * (98 – 25) = 7.665 J

98-58 Q = 4.2 * $10^3$ * 25 *$10^{-6}$ * (98 – 58) = 4.2 J

58-72 Q = 4.2 * $10^3$ * 25 *$10^{-6}$ * (72 – 58) = 1.47 J

72-98 Q = 4.2 * $10^3$ * 25 *$10^{-6}$ * (98 – 72) = 2.73 J

10 cycles $Q_{total}$ = 7.665 + (4.2+1.47) * 10+2.73 * 9 = 88.9 J

The thermocycling conditions of iDR was as follows: 30 min at 37℃.

25-37 Q = 4.2 * $10^3$ * 25 *$10^{-6}$ * (37 – 25) = 1.26 J

To highlight the difference, we've doubled the energy consumption.

### Section S5. Sequencing on an Illumina Hiseq 4000 platform.

**Sample collection and preparation.** DNA degradation and contamination were monitored on 2% agarose gels. DNA purity was checked using the NanoPhotometer spectrophotometer (IMPLEN, CA, USA). DNA concentration was measured using Qubit DNA Assay Kit in Qubit 2.0 Flurometer (Life Technologies, CA, USA).

**Library preparation for sequencing.** A total amount of 700 ng DNA per sample was used as input material for the DNA sample preparations. Sequencing libraries were generated using NEB Next® Ultra DNA Library Prep Kit for Illumina® (NEB, USA) following manufacturer’s recommendations and index codes were added to attribute sequences to each sample. Briefly, the Chip DNA was purified using AMPure XP system (Beckman Coulter, Beverly, USA). After adenylation of 3’ ends of DNA fragments, the NEB Next Adaptor with hairpin loop structure were ligated to prepare for hybridization. Then electrophoresis was used to select DNA fragments specified in length. 3 μL USER Enzyme (NEB, USA) was used with size-selected, adaptor-ligated DNA at 37°C for 15 min. At last, the products were purified (AMPure XP system) and library quality was assessed on the Agilent Bioanalyzer 2100 system.

**Clustering and sequencing.** The clustering of the index-coded samples was performed on a cBot Cluster Generation System using HiSeq 4000 PE Cluster Kit (Illumina) according to the manufacturer’s instructions. After cluster generation, the library preparations were sequenced on an Illumina Hiseq 4000 platform and 150bp paired-end reads were generated.

**Section S6. The bioinformatic statistical analysis.**

We stitched the reads pairs using PEAR used for oligo copy distribution, error and dropout ratio analysis.

The sequenced reads were aligned with the given sequences (synthesized by Twist Bioscience and CustomArray) by BLAST. Here, the reads without error containing substitution and indel were defined as M0G0 reads, and the reads with an error including substitution or indel and with two errors including substitution and indel were defined as M1G1 reads whose coverage, number and DNA sequences can be obtained via M0G0_M1G1.pl (fig. S3). The M0G0 and M1G1 reads were considered as valid reads.

The coverage and number could be achieved by Valid_Coverage_Number.pl (fig. S3). The frequency was achieved via the number dividing by total number of given sequences. Then the distribution of number of reads per each given sequence was displayed (Figs. 2F and 3D).

The distribution of the number of reads was displayed through analyzing coverage and number obtained by Obtain_Payload.pl (for obtaining valid DNA sequence namely payload), Cluster.pl and Sort.pl.

The coverage of aligned sequences was sorted from small to large and numbered them in sequence. Top 30% of the serial number was selected and the frequency of these reads was calculated (Fig. 2G).

115 sequences (1% * number of oligos in file) with top, bottom, and middle (mode value) coverage of #1 PCR and #1 iDR acted as a reference database. Per million valid sequences of both #5, #10 PCR and #5, #10 iDR were aligned with the chosen sequences. Then the percentage of reads was calculated via dividing by 1 million valid reads (fig. S6).

To depict the normality of oligo distribution, we used the piecewise function.

$$M = F(a + \triangle) - F(a - \triangle)$$

or

$$F(a) = \int_{a-\triangle}^{a+\triangle} f(x)\,dx$$

Here, F(x) is distribution function, f(x) is probability density function, a is the mean of oligo distribution, M is offset by defining △ as the distance (△ can be different values under the condition of △≤a). The area under the histogram is 1. In our study, △ is 50.

All sequenced reads were aligned with the actual reference sequences by basic sequence alignment program BLAST to screen out these reads with errors containing substitution, insertion, and deletion (henceforth referred to simply as “errors”) at the payload of individual sequences. And the number of reads with an error, two errors, three errors, ……, ten errors, more than ten errors in individual sequences were counted in detail by Mismatch_Analysis.pl and Gap_Analysis.pl and the frequency were calculated through the number of these reads dividing by the total number of noisy reads (fig. S4).

### Section S7. DNA decay caused by thermal condition.

Half-life of DNA extrapolate according to the Arrhenius Equation with activation energies of 155 ± 10 KJ mol$^{-1}$ and compared to literature data on DNA stability in solution previously reported. The temperature – logarithm of time curve of DNA decay was fitted as follows. In iDR, the optimal reaction temperature is 37℃. Under this condition, DNA decays within 9,927 h. However, DNA decays only within 2.6 h at 98℃，which is necessarily performed in PCR.

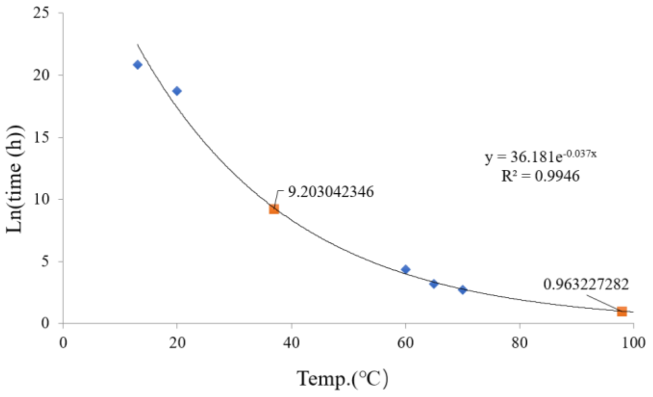

**Section S8. Minimum sequencing resource for perfect decoding.**

Taking pool 1 as an example. Oligo pool 1 is composed of 11520 oligos. We theoretically recovered the file at a coverage of 426 x in #10 PCR by fitting curve of the coverage-dropout ratio, which showed as follows. We successfully retrieved the file at a coverage of 12.5 x in #10 iDR. The percentage of valid reads are 48.58% and 92.24% among corresponding noisy reads of #10 PCR and #10 iDR respectively. Therefore, the total noisy reads which recovered the information with 100% accuracy required 11520*426/48.58% = 10,101,935 and 11520*12.5/92.24% = 156,115 of PCR and iDR separately. The ratio of noisy reads needed for successful decoding of #10 PCR to #10 iDR was 64-folds.

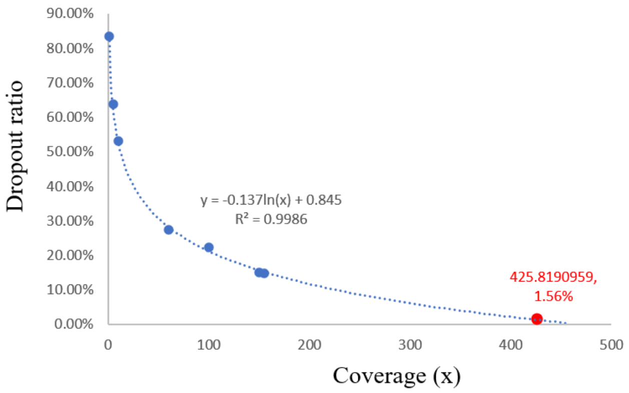

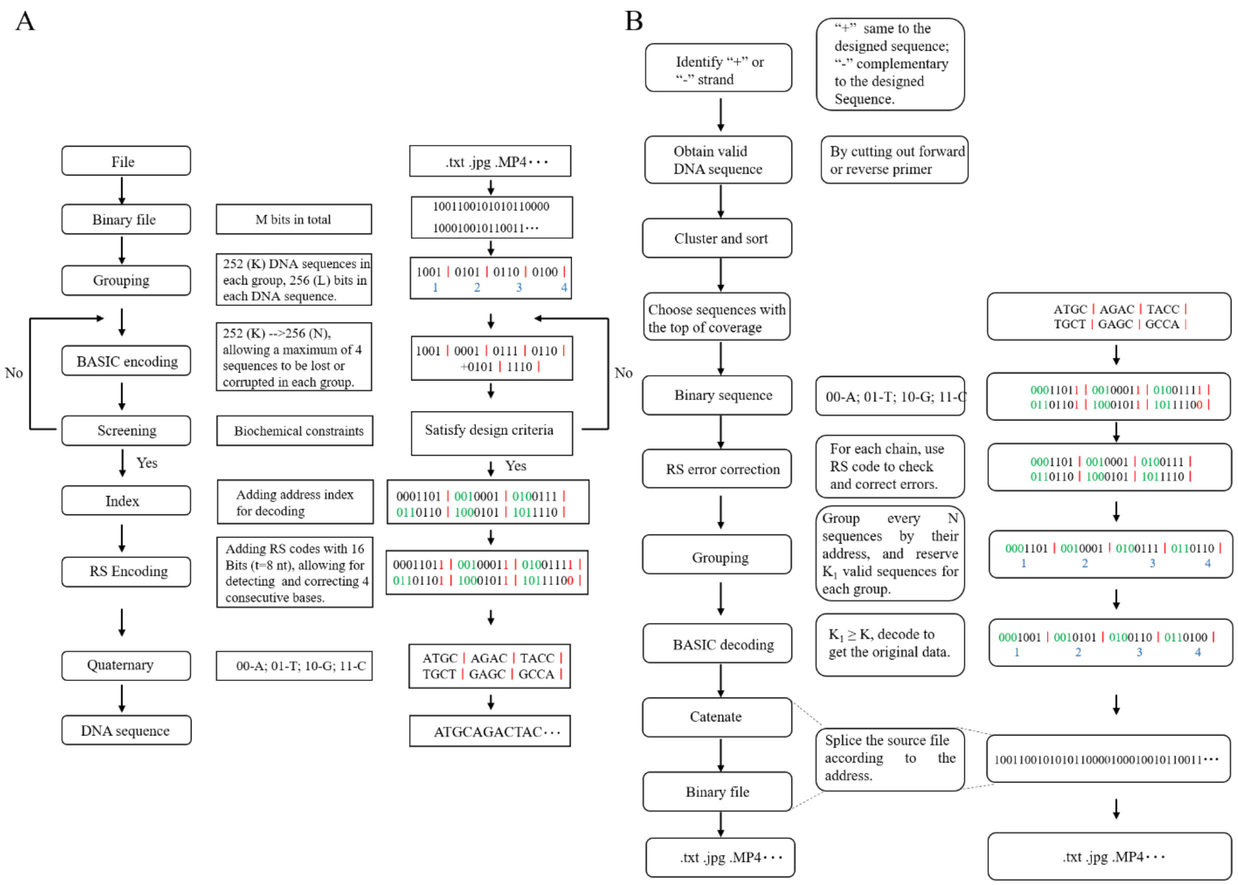


**Fig. S1. Workflow of DNA BASIC coding. (A)** (Left) The process of encoding in detail. K and L are arbitrarily defined. Here, K is the number of DNA sequence in each group; L is the length of binary data in each DNA sequence; K codewords of length L are encoded into N codewords by polynomial matrix operations on the polynomial cycle domain $\mathcal{C}_p$, where N>K, N-K is the maximum number of corrupted or lost sequences per group. Otherwise, we would not be perfect decoding. RS codes can detect and correct error within t/2 bases in each DNA sequence (Here, N=256, K=252, L=256, and t=8 in the design of pool 1 and pool 2; N=256, K=252, L=192, and t=8 in the design of pool 3). (Right) Example file. **(B)** DNA BASIC decoding. (Left) The process of decoding in detail, which mainly contain cluster and sort of DNA sequence, RS error correction, and BASIC decoding. (Right) Example file.

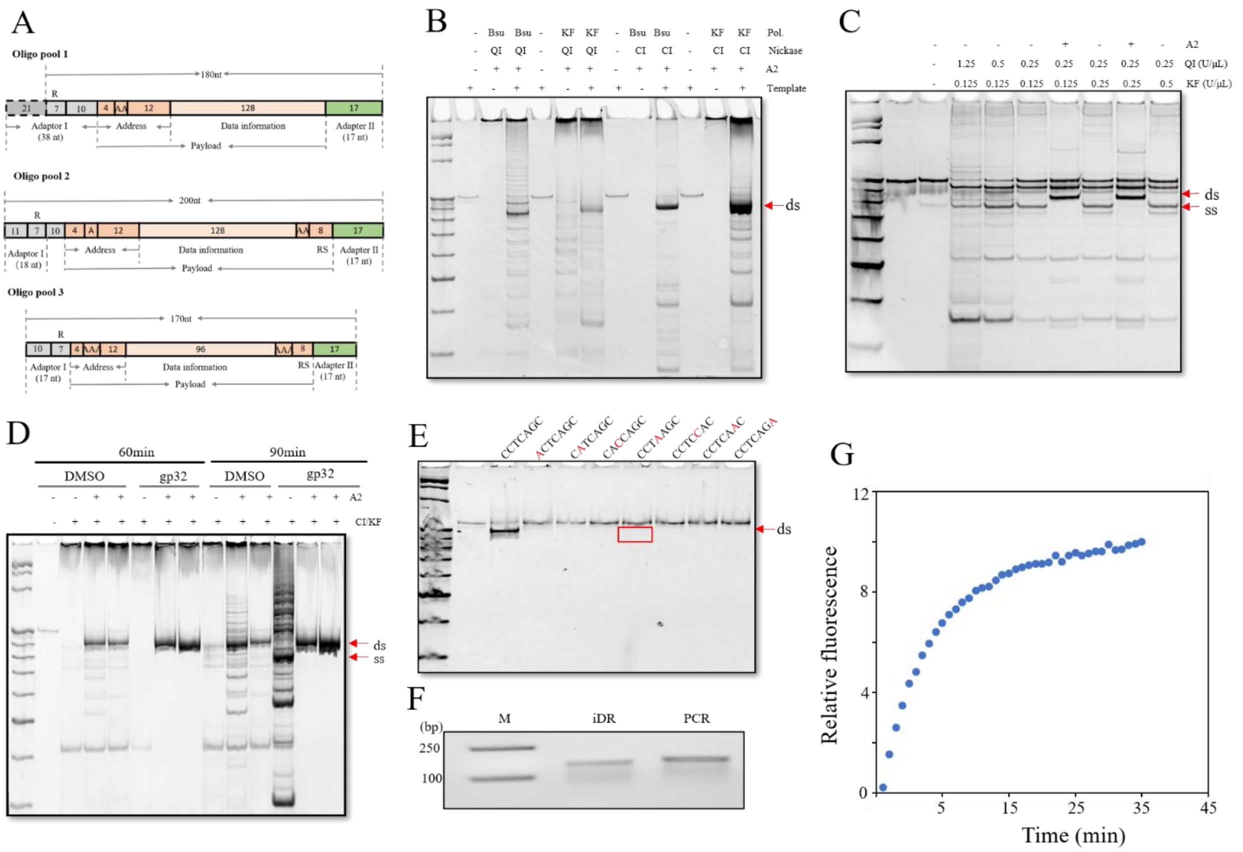


**Fig. S2. Characterization of iDR reaction. (A)** The structure of oligos. 11,520 oligos (Oligo pool 1, Twist Bioscience) without RS code were encoded and synthesized. And 11,520 oligos (Oligo pool 2, Twist Bioscience) with RS code were encoded and synthesized. 86,528 oligos (Oligo pool 3, CustomArray) were encoded and synthesized. (R: recognition sequence of nickase). **(B)** iDR reaction under 37°C. The combination of nickases and polymerases were Nt.BspQI/Nt.BbvCI and Bsu DNA polymerase/Klenow fragment (3’→5’ exo$^{-}$). **(C)** The concentration of QI and KF was optimized. The yield of iDR product was increased with increasing amount of QI rather than KF. **(D)** Additives was also optimized. The yield of iDR product was increased while the by-product was decreased with addition of T4 Gene 32 protein (gp32). **(E)** The iDR product was observed when CCTCAGC, recognition sequence of Nt.BbvCI, was converted into CCTAAGC. It demonstrated that Nt.BbvCI was unspecific to recognize CCTCAGC. Thus, CCTNAGC was evaded in the sequence we encoded. **(F)** 2% agarose results of iDR and PCR products. **(G)** Real-time iDR. The iDR reaction mixtures contained 1 μL of the template (10 ng/μL), 0.25 mM dNTPs, 2.5 μL 10x NEBuffer 2, 0.08 U/μL Nt.BbvCI, 0.16 U/μL KF polymerase (exo$^{-}$), 4 μM T4 Gene 32 protein (gp32), 0.2 mg/mL BSA, 0.5 μM adaptor 2 (For production of ssDNA, adaptor was not added.) and 0.5 μL 3.75xSYBR Green I. The mixtures were incubated at 37℃ for 30 min and detected every 30s.

Note: Red arrow represents the product of iDR. QI: Nt.BspQI; CI: Nt.BbvCI; Bsu: Bsu DNA polymerase, large fragement (3'→5' exo$^{-}$); Klenow fragment (3'→5' exo-). A2: Adaptor 2.

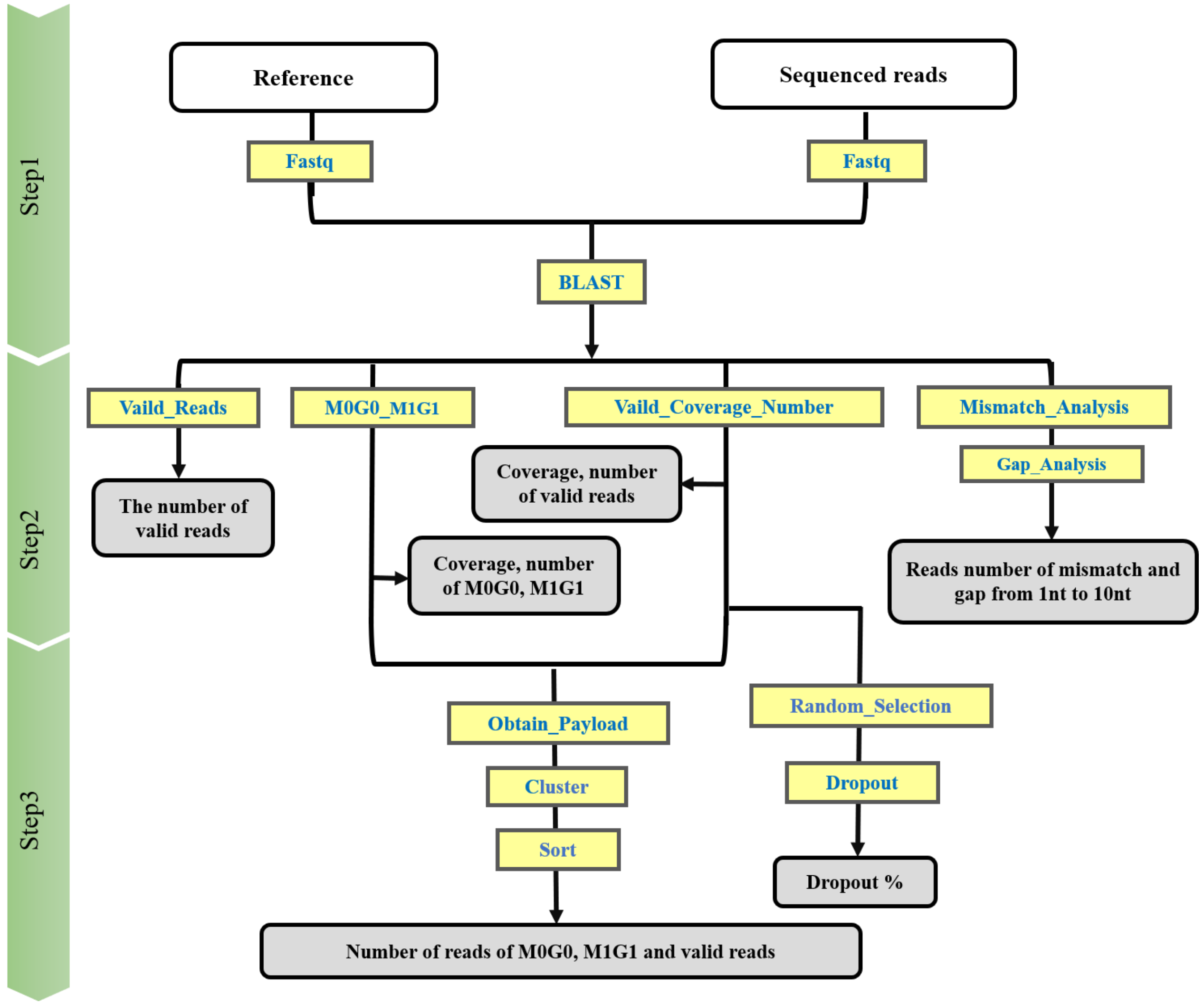


**Fig. S3. The workflow of bioinformatic statistical analysis.** Bioinformatics analysis programs are in the yellow boxes. Results generated by bioinformatics analysis programs are gray boxes.

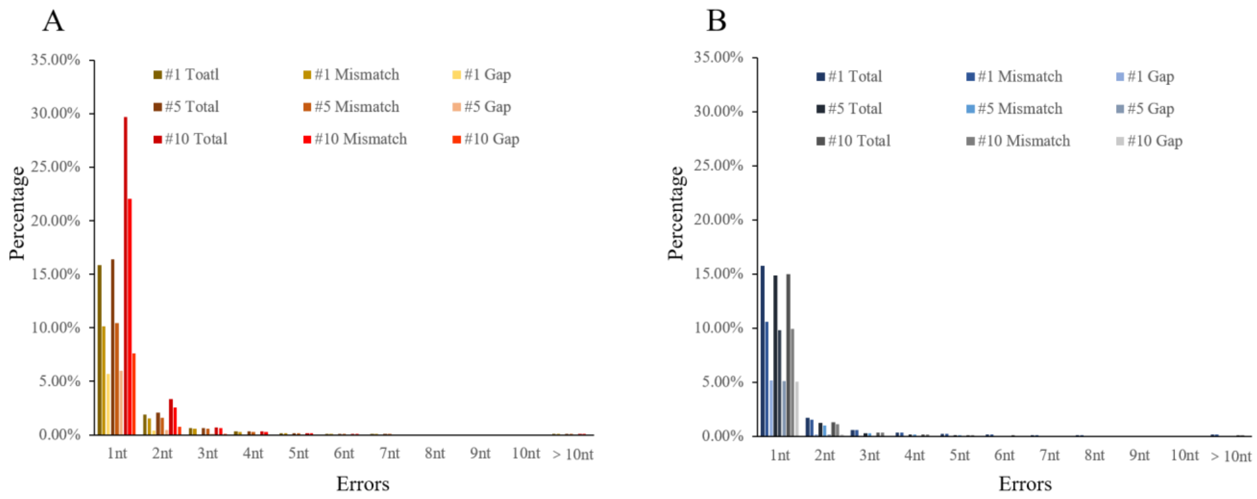


**Fig. S4. Characterization of error distribution. (A)** The percentage of the reads with errors containing substitution and indel among total reads respectively in repetitive #1, #5, #10 PCR (Pool 1, Twist Bioscience). **(B)** The percentage of the reads with errors containing substitution and indel among total reads respectively in #1, #5, #10 iDR (Pool 1, Twist Bioscience).

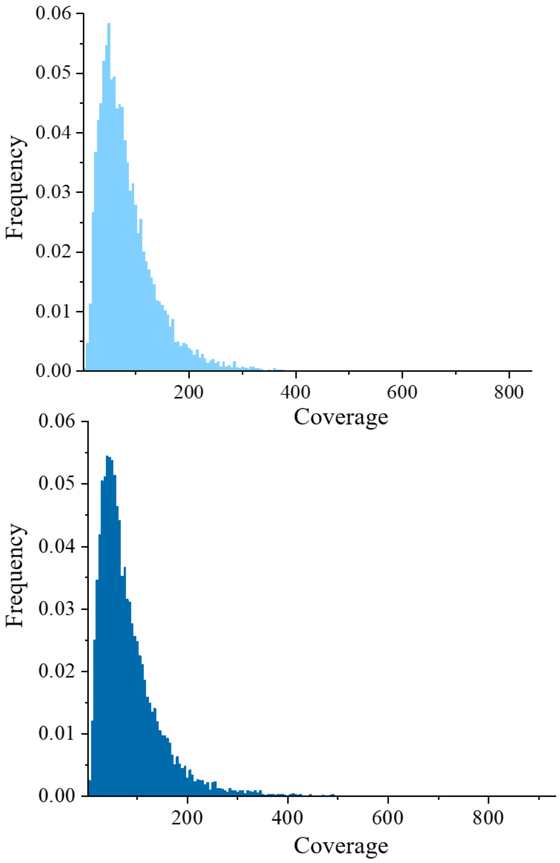


**Fig. S5. Characterization of effect of DNA oligo physical state.** The distribution of the number of reads per each given sequence of free-iDR (without Streptavidin magnetic beads, the upper portion) and iDR-amplified (the lower part) the oligo pool (Pool 1, Twist Bioscience).

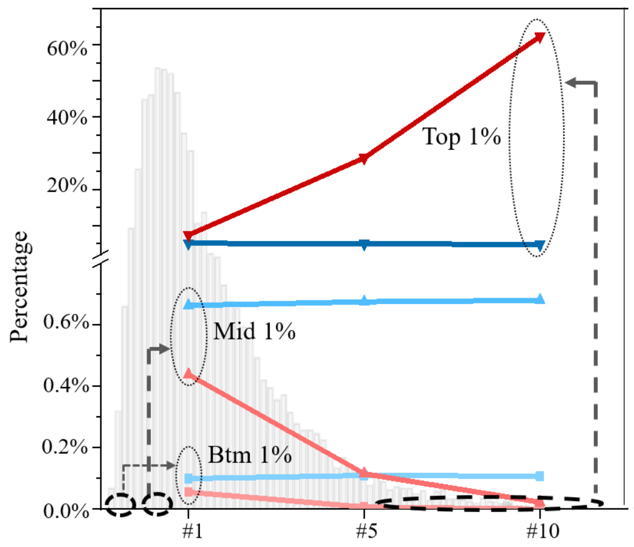


**Fig. S6. Quantification of oligo spreading with amplification.** The percentage of some certain sequences (1% of total number) with the bottom, middle and top coverage in PCR#1 and iDR#1 was investigated in ten serial PCR and iDR separately (Section S6). The frequency of 116 oligos (1% of the total number, dark red) with the top coverage rose to 62.3% (#10 PCR), compared with 7.4% (#1 PCR). But, the frequency of 116 strands (1% of the total number) with the middle (Pink) and bottom (light pink) coverage was down to 0.011% and 0.0002% separately in #10 PCR. However, the frequency of 116 oligos whatever the coverage was had remained stable in iDR (blue-colored items).

**Table S1. Digital files encoded.** Files encoded within these 2.85 MB of data. Both pool 1 and pool 2 contain 11,520 DNA sequences. Pool 3 contains 86,528 DNA sequences.

| Data | File Size | Number of DNA strands | Pool |
|---|---|---|---|
| Central dogma (.jpg) | 35 KB | 990 | Pool 1/2 |
| DNA helix (.gif) | 81 KB | 2292 | Pool 1/2 |
| China Classical literature (.txt) | 164 KB | 4641 | Pool 1/2 |
| A Brief History of Element (.txt) | 34 KB | 962 | Pool 1/2 |
| Panda burn incense (.rar) | 66 KB | 1867 | Pool 1/2 |
| Human Mitochondrial | 65 KB | 768 | Pool 1/2 |
| Bitcoin (.txt) | 165 KB | 7019 | Pool 3 |
| Dictionary of idioms (.txt) | 70 KB | 2978 | Pool 3 |
| Black hole (.jpg) | 86 KB | 3659 | Pool 3 |
| Chaplin (.MP4) | 659 KB | 28034 | Pool 3 |
| JCVI-syn.0.1 | 1054 KB | 44838 | Pool 3 |
| **Total** | **2924 KB** | **109,568** | |

**Table S2. Decoding of digital file from DNA pool.** Samples with PCR or iDR amplified were whether could be successfully decoded under the condition of total sequenced reads (noisy reads) used. And the coverage was showed when the dropout rate was 1.56%.   Note: '*' Total noisy reads with missing 14.1% of given sequences (11,520) could not decode original file. '**’ The dropout rate was beyond 1.56% even when total noisy reads were used.

| Pool | Sample | Perfect Decoding (Noisy reads) | Coverage (Dropout=1.56 %) |
|---|---|---|---|
| Pool 1 | Free-iDR | Yes | 9x |
| | PCR | Yes | 12.5x |
| | ss | Yse | 18x |
| | #1 PCR | Yes | 17x |
| | #5 PCR | Yes | 160x |
| | #10 PCR | No* | -** |
| | #1 iDR | Yes | 11x |
| | #5 iDR | Yes | 12x |
| | #10 iDR | Yes | 12.5x |
| Pool 2 | PCR | Yes | 6x |
| | iDR | Yes | 5x |
| Pool 3 | PCR | Yes | 65x |
| | iDR | Yes | 49x |

**Table S3. Primer list.** Primer sequences used. Note: The recognition sequences of nickase was marked with bold fonts.

| Name | Sequence (5’→3’) | For oligo pool |
|---|---|---|
| Adaptor I | GTCCCGCTCATGCATCACCTA**CCTCAGC**TCAACTCACT | |
| Adaptor I-1 | GTCCCGCTCATGCATCA | Pool 1 |
| Adaptor II | TCCACGACGATCAGACT | |
| Adaptor II-1 | Biotin-AAAAATCCACGACGATCAGACT | |
| probe | FAM-AGATCAATTAATACGATACCTGCGTTT | |
| splint | CTCGGAAGAGCTGAAAACGCAGGTATCG | |
| T1 | GTCGCTAACAGAGTAACCTCCTCAGCTCTTCCGAGTCGGC<br>AGCACTGCATAATTCTCTTACTGTCATGCCATCCGTAAGAT<br>GCTTTCTGTGACTGGTGAGTACTCAACCAAGTCATTCTGAG<br>AATAGTGTATGCGGCGACCGAGTTGCTCTTGCCCGGCGTC<br>AATACGGGATAATACCGCGCCACATAGCAGAACTTATGAG<br>TGGAGGTGTAAAGTG | Ligation |
| Adaptor I | TGCATCACCTA**CCTCAGC** | |
| Adaptor II | TCCACGACGATCAGACT | Pool 2 |
| Adaptor II-1 | Biotin-AAAAATCCACGACGATCAGACT | |
| Adaptor I | GTCGCCTTCT**CCTCAGC** | |
| Adaptor II | AGCGCTTTAAGCCAACA | Pool 3 |

**Table S4. Sequences were avoided in the process of encoding.**

| Serial number | Sequences |
|---|---|
| 1 | GCTCTTC |
| 2 | GAAGAGC |
| 3 | GCTGAGG |
| 4 | CCTCAGC |
| 5 | CCTAAGC |
| 6 | GCTTAGG |
| 7 | CCTTAGC |
| 8 | GCTAAGG |
| 9 | CCTGAGC |
| 10 | GCTCAGG |
| 11 | CCTCAGT |
| 12 | ACTGAGG |
| 13 | CCTCAGG |
| 14 | AGATAG |
| 15 | TGTTGG |
| 16 | GAGCTG |
| 17 | AGTCTG |
